# Learning to operate an imagined speech Brain-Computer Interface involves the spatial and frequency tuning of neural activity

**DOI:** 10.1101/2023.09.11.557181

**Authors:** Kinkini Bhadra, Anne Lise Giraud, Silvia Marchesotti

## Abstract

Brain-Computer Interfaces (BCI) will revolutionize the way people with impaired speech production can communicate. While recent studies confirm the possibility of decoding imagined speech based on pre-recorded intracranial neurophysiological signals, current efforts focus on collecting vast amounts of data to train classifiers, rather than exploring how the individual’s brain adapts to improve BCI control, an important aspect given the known problem of “BCI illiteracy”, the inability of some individuals to operate a BCI. This issue can be investigated by providing real-time feedback to allow users to identify the best control strategy. In this study, we trained 15 healthy participants to operate a simple binary BCI system based on electroencephalography (EEG) signals through syllable imagery for five consecutive days. We explored whether BCI-control improves with training and characterized the underlying neural dynamics, both in terms of EEG power changes and of the neural features contributing to real-time classification. Despite considerable interindividual variability in performance and learning, a significant improvement in BCI control was observed from day 1 to 5. Performance improvement was associated with a global EEG power increase in frontal theta and a focal increase in temporal low-gamma, showing that learning to operate an imagined-speech BCI involves global and local dynamical changes involving low- and high-frequency neural features, respectively. These findings indicate that both machine and human learning must be considered to reach optimal controllability of imagined-speech BCI, and that non-invasive BCI-learning can help predict the individual benefit from an invasive speech BCI and guide both the electrode implantation and decoding strategies.

## Introduction

Neurological disorders of language such as aphasia, amyotrophic lateral sclerosis and locked-in syndrome can disrupt natural speech resulting in a reduced quality of life for both patients and caregivers ^1,2^. One promising approach to restore language communication is to decode imagined speech directly from neurophysiological signals through a brain-computer interface (BCI) and translate them into text or synthesized speech. This approach has raised two important challenges: how the machine can decode neural signals, and how the patient can optimize its interaction with the decoder. For the latter, providing a feedback to the user in real-time is crucial.

Recent years have seen great advances in the field of speech-BCIs, most often through the decoding of motor representations of vocal tract movements from intracranial electrophysiological recordings ^3–8^, that have allowed reaching a decoding speed of 78 ^4^ words per minute. Such an approach, however, will not be suitable for disorders of language where speech production areas are damaged, such as in post-stroke aphasia. A BCI appropriate for these disorders would require decoding representation of speech units produced through imagined, rather than attempted speech, in particular involving the language temporo-frontal system. Also named covert speech, imagined speech consists in the internal pronunciation without self-generated audible output ^9,10^, thus without the involvement of the musculoskeletal system. In patients with aphasia, both subjective and objective assessments indicate that imagined speech is better preserved than spoken language ^11–13^, even in presence of severe overt (i.e. articulated and audible) speech deficits ^14^. Thus, investigating imagined speech can have important implications for patients with aphasia, while remaining a valid approach for other disorders of speech production.

Although previous studies have characterized the neural correlates of imagined speech ^15^, most often in comparison with overt speech ^16–21^, only a handful of BCI studies have attempted to decode imagined speech in real time, with promising but often limited effectiveness ^22–25^. This is due to different challenges and limitations primarily pertaining to the weakness of imagined speech signals as compared to overt speech ^16,17,19,20,26^, the difficulty to precisely identify the onset of speech imagery ^26^, inter-individual differences in the ability to control the BCI ^27,28^, and to the technique employed to record brain activity.

The state-of-the-art approach is the use of intracranial recordings such as electrocorticography (ECoG) and stereotactic EEG (sEEG), which allow neural sampling from key language regions with higher spatial resolution, and the possibility to use high-frequency neural activity. Exploiting these experimental advantages, imagined speech decoding for BCI-control has been first attempted using ECoG to decode imagined phoneme pronunciation versus rest from the perisylvian area ^22^. Although decoding accuracy was highly above chance, this study did not provide evidence that the method could be used to discriminate between two imagined speech units in real-time. More than a decade later, sEEG was used to synthesize imagined speech in real-time into a continuous acoustic feedback from high-gamma activity in the frontal cortex and motor areas ^24^. Although reconstructed speech was unintelligible and less accurate for imagined than overt speech, this study was the first proof-of-concept that imagined speech could be used for naturalistic communication with a speech neuroprosthesis. More recently, impressive real-time control (up to 91%) was achieved by decoding eight imagined words from single-neuron activity in the supramarginal gyrus ^25^, highlighting the superior effectiveness of decoding speech from individual neurons. Yet, using intracortical recording for speech-BCI remains a clinical and ethical challenge, owing to the high risk of clinical complications (loss of contacts, infection) leading to possible explantation and the loss of the new communication means, a dramatic outcome for the patient. Much research and clinical efforts are still required to optimize the success of future intracortical speech-BCI.

Capitalizing on its far greater accessibility, several studies have employed surface EEG for decoding offline (i.e. open-loop) a wide variety of speech units imagery, most often in binary classification paradigm ^9,29^ (see for reviews ^9,30,31^). However, nearly all studies address imagined speech decoding from an engineering perspective, their main goal being the optimization of current classifiers to boost decoding accuracy (see ^32^ for a review of classification methods. Despite the great amount of data samples used in offline decoding as compared to real-time control (i.e. online, closed-loop), and the possibility of applying computationally demanding decoders, offline performance from pre-recorded dataset remain below 80% when discriminating between two imagined speech units and around 60% for a three-class problem ^29,33–36^.

In the single BCI study that used EEG for real-time speech imagery decoding ^23^, performance remained below 70% in discriminating between “yes” and “no”. Interestingly, however, this study pointed out important inter-individual differences in BCI-control, with accuracies varying between 53.75% and 95% ^23^.The variability in control abilities is well known in motor-imagery EEG-BCIs, in which up to 50% of participants are unable to achieve sufficient BCI-control ^37,38^. Given the lack of speech-imagery EEG-BCI studies and the fact that invasive-BCI studies are mostly single-case, it remains to be assessed whether speech-BCI skills can be improved with training. In the present study, we addressed speech-BCI controllability from a neurophysiological rather than neuroengineering side. We addressed whether BCI-control performance can be trained together with exploring interindividual variability and the neural mechanisms underpinning the acquisition of these new skills. We designed a closed-loop BCI system based on electroencephalography signals (EEG) to decode in real-time the imagery of two syllables /fɔ/ and /gi/, chosen for their contrasted phonetic features and trained 15 healthy participants to control the BCI for 5 consecutive days. This study thus addresses both the variability and the dynamic range that can be achieved via training a whole brain EEG speech-imagery BCI.

## Materials and Methods

### Participants

Fifteen healthy participants (5 females, mean age 23.9 years, SD ±2.3, range 19-29) took part in this study which was approved by the local Ethics Committee (Commission cantonale d’éthique de la recherche, project 2022-00451) and was performed in accordance with the Declaration of Helsinki. All participants provided written informed consent and received financial compensation for their participation. All participants were right-handed.

### Experimental paradigm

Participants took part in the study daily for 5 consecutive days, at the same time of the day. To avoid potential effects of individuals’ circadian rhythm, each participant began the training at the same time each day. Each session lasted approximately 2.5 hours, amounting to a total duration of 12-13 hours of experimental time per participant. The experiment took place in an optically, acoustically, and electrically shielded room.

#### Syllable imagery

During the daily BCI training, participants were asked to imagine saying one of the two syllables /fɔ/ and /gi/, chosen for their contrasted phonetic features regarding consonant manner (fricative vs plosive), place of articulation (labiodental vs velar), vowel place (mid back vs high front) and rounding (rounded vs unrounded). This choice was motivated by previous results showing differences in the neural responses associated with these different phonetic features ^20,39–41^ and was expected to maximize the discriminability between the EEG signals associated with the imagery of each syllable, thereby facilitating their decoding by a classifier.

Participants were instructed to imagine saying each of the two syllables, focusing on the kinesthetic sensation they would experience if they would pronounce the syllable, rather than imagine hearing oneself speaking or imagine the syllable written in characters. They were explicitly told to avoid any movements during the imagery, especially those involving the face, not to mouth nor whisper. They were informed that the muscular activity of the face was being monitored with EMG electrodes throughout the entire experiment to control for such movements.

### EEG acquisition and BCI loop

#### EEG recording

Neural data were recorded using a 64-channel ANT Neuro system (*eego mylab*, ANT Neuro, Hengelo, Netherlands) at a sampling rate of 512 Hz using electrode AFz as ground and CPz as reference. Channels’ impedance were kept below 20 kΩ throughout the experiment. Electromyography signals (EMG) were recorded from two facial muscles on the right side of the participant’s face, the zygomaticus major and the orbicularis oris, to control for potential articulatory muscles’ activation despite our explicit instructions to avoid any movement ^42^. The placement of EMG electrodes on the right side was determined by the participants’ handedness (all right-handed), as the dominant side of the face typically matches the dominant hand, and therefore tends to exhibit more pronounced movements during speech production^43,44^. EEG and EMG data were acquired using Lab Streaming Layer (LSL, https://github.com/sccn/labstreaminglayer).

During the EEG recording and while operating the BCI, participants sat comfortably on a chair in front of a computer screen while keeping their hands on their thighs. They were instructed to avoid any possible physical movements, especially eye blinks and mouth articulation, while performing the speech-imagery.

#### BCI loop

The EEG-BCI loop was developed using an adapted version of the framework *Neurodecode* (Fondation Campus Biotech Geneva, https://github.com/fcbg-hnp/NeuroDecode), already used in previous BCI studies ^45,46^. On each training day, the BCI-control included two sessions, *an offline session* in which data were recorded for the classifier’s calibration and an *online* part where participants controlled a visual feedback in real-time. Therefore, the classifier was different on each experimental day. Participants were asked to use the same imagery strategy throughout the entire duration of the BCI training (see *Syllable imagery* section).

#### Offline session and classifier calibration

EEG data were acquired while participants performed syllable imagery without receiving any real-time feedback (offline runs) and subsequently used to calibrate the classifier. Each offline trial began with a text indicating the trial number (1 s), followed by a fixation cross (2 s), and a written cue indicating which of the two syllables participants had to imagine pronouncing (2 s). After the cue disappeared, an empty battery appeared on the screen, which then progressively filled for 5 s (Fig. 1a). Participants were instructed to start imagining pronouncing the syllable immediately after the battery appeared on the screen and stop when the battery was filled. They were explicitly told that the battery filling was independent of their brain signals. At the end of the 5 s imagery period, the battery was displayed as it appeared at the last filling level, for 2a additional seconds, with the tip of the battery turning yellow to indicate the participant to stop imagining. Participants had 5 s to rest while the text ‘Rest’ was displayed on the screen.

**Fig. 1.**
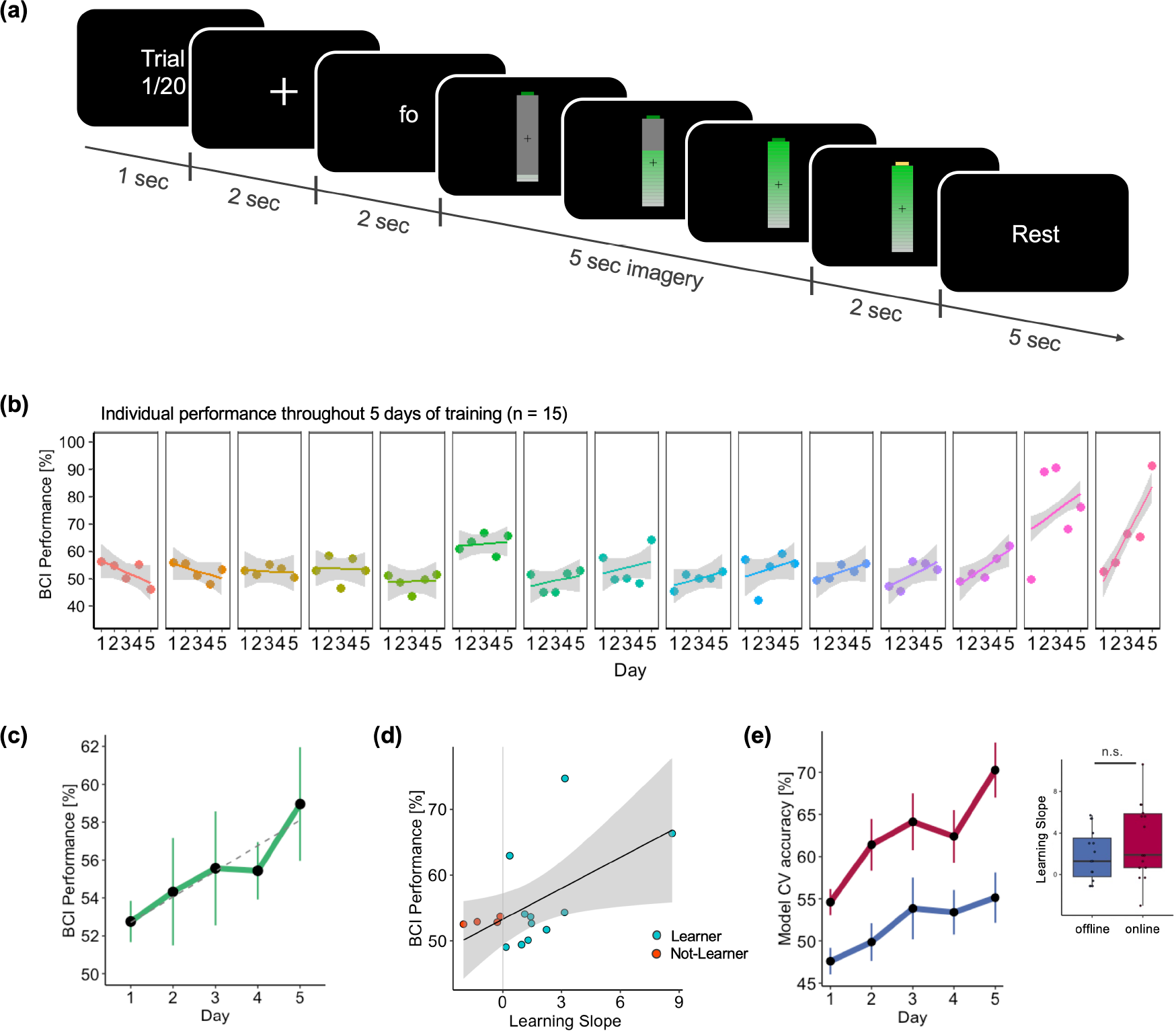
Experimental paradigm and online BCI performance: **(a)** Experiment Paradigm. Each trial began with the presentation of the trial number, followed by a fixation cross. The last second of the fixation cross was considered as a baseline for the EEG analysis. Next, participants were instructed to imagine pronouncing either the syllable /fɔ/ or /gi/ as indicated by a text on the screen. They had to continue performing the imagery throughout 5 s in the offline session, or until the battery was filled during the real-time control (online session). To indicate the end of the imagery period, the battery appeared full and the top turned yellow. **(b)** Online BCI performance (%) for each participant ordered according to the value of the learning slope (from lowest to highest). Dots represent the average BCI-control performance for each training day, with the corresponding regression line. **(c)** Average BCI online performance (%) over the 5 training days. Dots represent the average performance across participants with error bars indicating the standard error of the mean, and the significant linear regression (dashed line). **(d)** Correlation between individual learning slopes and average BCI performance across the 5 training days. Positive and negative learning slopes are indicated by orange (learners) and cyan (not-learners) dots, respectively. **(e)** Cross-validation (CV) accuracy obtained by computing the classifier in the offline (blue) and online (red) sessions on each day (left). The box plot (right) shows the learning slopes obtained by fitting a linear model per participant and session (offline in blue and online in red) using CV accuracies on each training day. There was no significant difference between the two sessions.

There were a total of 40 trials per syllable, arranged in 4 blocks each consisting of 20 trials per syllable, with short breaks in between blocks. The *offline session* lasted approximately 25-30 min.

Offline data was then used to *calibrate the decoder*. Features were extracted by computing the power spectral density (PSD) of the EEG signal from 1 to 70 Hz (with a 2 Hz resolution) using a sliding window of 500 ms and 20 ms overlap. The PSD was calculated for each EEG channel excluding the three electrodes placed over the mastoid region bilaterally and the reference channel, leading to 61 channels. Therefore, there were a total of 2135 features (61 channels and 35 frequencies), each of which consisting of a channel-frequency pair. These features were then fed to a random forest (RF) algorithm to extract the classifier parameters (i.e. the covariance matrix). This nonlinear classifier has already been proven effective in previous two-class BCI studies ^47^ and is known to be robust to overfitting ^48^. The RF classifier assigns a weight (expressed in percentage) to each feature, indicating its relative contribution to the classification. A 8-fold cross-validation was performed to test the model validity and calculate the offline classification accuracy. Beside being used for real-time control, we analysed features’ weights to investigate which brain regions and frequency bands contribute the most in classification and identify changes across days (see *Data Analysis* section).

#### Online BCI-control

During the online part of the experiment, the RF classifier was applied in real-time to the EEG data to decode which of the two syllables the participant was imagining, and accordingly, provided a real-time feedback to the user. Trials were the same as during the offline part, except that this time the battery could fill or empty based on the output of the classifier, hence informing the participants about their neural performance. Participants were instructed to fill the battery by performing the same imagery task as before.The mapping of the decoder output to the battery feedback at each time sample was done in such a way that if the probability output by the classifier matched the cued syllable, the battery would fill, otherwise it would empty. The real-time control continued either until the battery was full or until a 5 s timeout. As for the *offline* session, there were 40 trials per syllable, divided into 4 blocks.

To boost participants’ motivation and keep them engaged in the task throughout the entire training period, we provided monetary bonuses based on performance. Participants received an additional 10 CHF for each experimental day in which they performed above chance (50%) and was higher than on the previous day.

### Data Analysis

#### BCI Performance and CV accuracy

Participants’ performance in controlling the BCI (*BCI-control performance*) was calculated by considering, for each trial, the rate (%) of trials where the classifier’s output corresponded to the cued syllable. To investigate whether training resulted in an increase in BCI-control performance, we performed a planned contrast analysis, considering the average performance during each training days for each participant, and testing for a linear increase or decrease from day 1 to 5 (numeric contrast using as weights −2,−1,0,1, and 2, respectively for day 1 to 5). This set of contrasts was tested in a linear mixed model (LMM), with participants as a random factor. The same statistical approach was used to probe changes across days in other dependent variables (e.g. features weight and power modulation evolution), and it is hereinafter referred to as “*LMM with planned contrast*”.

We investigated whether the learning dynamics was related to the individual’s ability to control the BCI. To do so, we fitted, separately for each participant, a linear model considering the average performance on each day and extracted the individual learning slope. We then computed the Pearson correlation between the average performance during the entire training period and the learning slope.

Additionally, we tested for differences in performance between the two syllables (2-tailed paired t test) and between blocks (one-way repeated measures ANOVA with *Block* number as within-participant factor, and 2-tailed paired t tests for post hoc comparisons).

To further assess potential learning mechanisms across training, we analysed the open-loop decoding accuracy across the training period as a proxy of BCI performance. For this, we considered the cross validation (CV) accuracy of the RF model computed to calibrate the classifier using the *offline* data (see “*classifier calibration*” section) and applied the same method to compute CV accuracy based on the *online* data. This approach pools together all data from an individual session (*offline* or *online*) and is thus different from the method used to compute the *BCI-control performance,* which is based on individual samples at the single-trial level. We tested for a linear increase in CV accuracy using the LMM with planned contrast, separately for the *offline* and *online* sessions. We compared the CV accuracy between the two sessions, averaged across days, with a 2-tailed paired t test between the CV-accuracies

To test for differences in learning between offline and online sessions, we extracted the learning slope by fitting, separately for each participant and each session (*offline*/*online*), a linear model considering the CV accuracy across the 5 days of training, using the same method as for the *BCI-control performance*. We then assessed differences in learning slope between off- and on-line sessions by performing a 2-tailed paired t-test.

#### Classifier features

Next, we investigated which brain regions and frequency bands contribute most to BCI-control and study changes in decoding patterns across training days. For this, we considered the feature weights obtained by training the RF classifier with the data acquired during the *offline* session. Each feature refers to a specific channel-frequency pair leading to a total of 61 (channels) x 35 (frequencies) features. The weight of each feature is expressed as a percentage, where higher values indicate a stronger contribution in discriminating between the two syllables.

First, we addressed whether better BCI-control is associated with higher features’ weights, reflecting a better discriminability between the two classes. To do so, we considered, separately for each participant, the sum of the weights across the first 200 features (i.e. irrespective of frequency or channel location, ranked according to their weight). This sub-sampling was necessary since the weight of each feature is expressed in percentage, thus the sum of all features would have led to the same value of 100%, preventing testing for difference across participants. The choice of this subset size was motivated by the fact that the cumulative sum of the first 200 features weights exceeded on average 50% (Supplementary Fig. 1a). We performed a Pearson correlation coefficient between the feature’s weight sum, averaged across days, and the average individual BCI-control performance (obtained as described in *BCI Performance and CV accuracy* section).

Next, we investigated the topography of the most discriminant features, as well as their frequency distribution. We considered the average weight over the 5 training days (1) separately for each individual frequency i.e. from 2 Hz to 70 Hz with a 2 Hz resolution disregarding to which electrode the features belonged and (2) across the scalp separately for each frequency value and frequency band, to obtain a topographical representation of the feature weights.

#### Features evolution over training

Subsequently, we quantified the evolution of the features weight over the five training days. First, to investigate global changes in the weights across training, we considered the weight sum of the first 200 features for each training day and ran the LMM with planned contrast analysis, testing for a linear change in the weight with training. A positive relationship between BCI-control performance and weight over the 5 days would reflect a behavioral improvement.

Next, we investigated changes due to training more specifically at the level of the decoding frequencies and brain regions. We first inspected changes by visualizing feature pairs as individual elements in a 2D map, with frequency values (from 2 Hz to 70 Hz, with 2 Hz resolution) and individual channels respectively represented on the x- and y-axis.

We quantified changes over time separately in the frequency and spatial domain (i.e. at the topographic level). To limit the number of multiple comparisons, we guided the spatial analyses by the results obtained in the frequency domain.

To identify frequency-wise changes to the discrimination between the two syllables, we considered the average weight across all features for each individual frequency value. We performed a “LMM with planned contrast” for each frequency value separately and assessed the statistical significance of the linear change across the 5 days of training with 1000 permutations, assigning randomly the day to which data belonged, within participants. Based on these results, we defined frequency intervals, over which performing the analysis in the spatial domain. We then considered the average weight across all features and ran a “LMM with planned contrast” for each electrode and each interval. The statistical significance was assessed with 1000 permutations, again by randomizing the factor *Day* on a single participant basis.

Next, we evaluated the relationship between changes in BCI-control performance across days and changes in the features space. To quantify the global feature changes in a single index per participant, we considered the euclidean distance between the feature weights of two consecutive days according to the following formula:

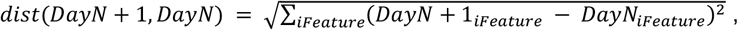

where N indicates the experimental Day and iFeature a Frequency-Channel pair (Fig. 2f). The four distance matrices (one for each couple of consecutive days) were then averaged to obtain an individual index per participant. Last, we correlated this index with the average BCI performance computed considering all training days.

**Fig. 2.**
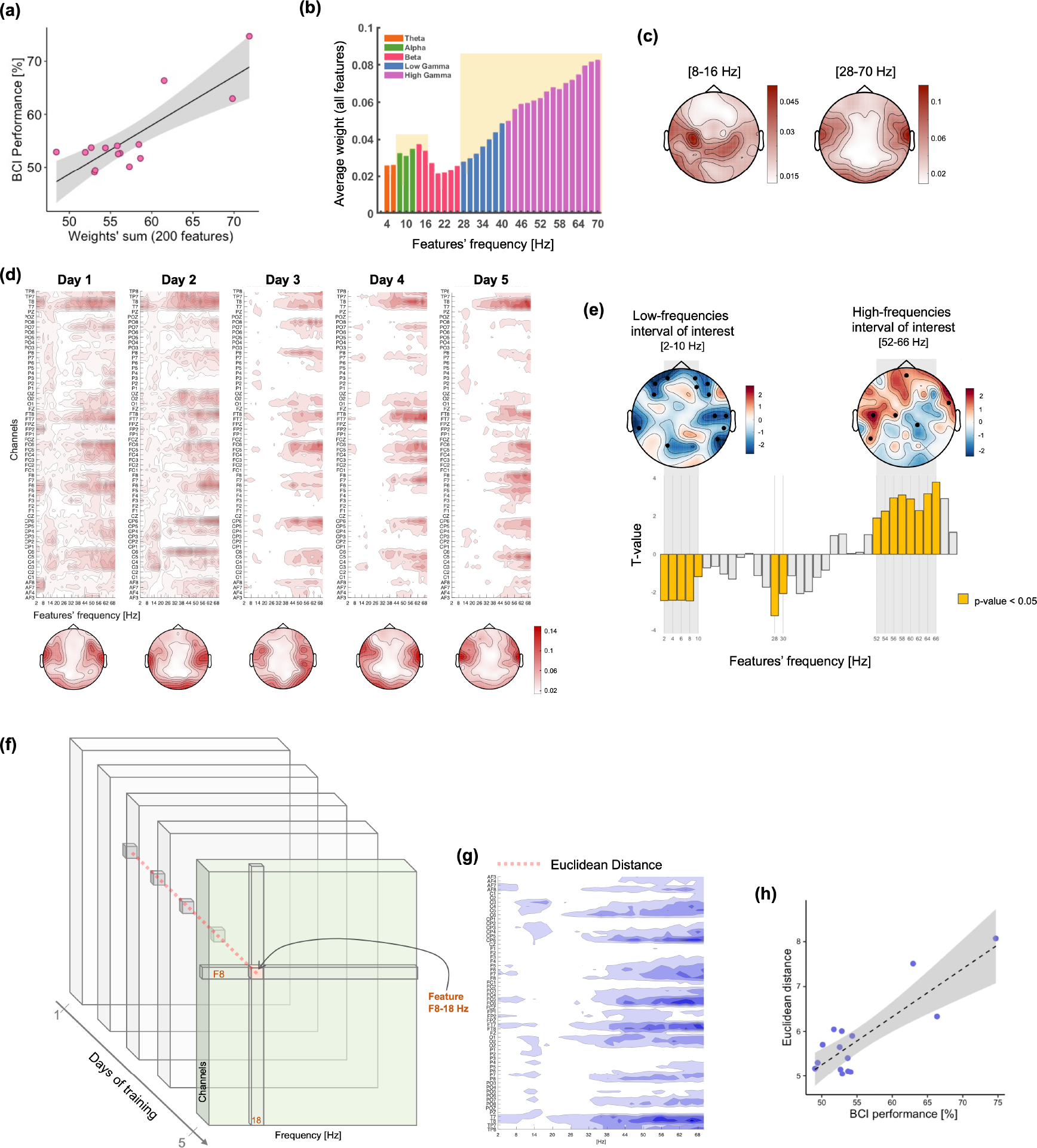
Random forest classifier features and their evolution over training. **(a)** Relationship between individual BCI-control performance and feature weights, obtained by considering the sum of the first 200 features, averaged across the 5 training days. **(b)** Average of features’ weights for each frequency across all experimental days. Colors indicate the different frequency bands (theta: orange, alpha: green, beta: pink, low-gamma: blue, high-gamma: magenta). The highlighted intervals indicate two frequency ranges with prominent contribution to syllable decoding: 8-16 Hz and 28-70Hz. **(c)** Corresponding topographies of the average weights for these two frequency intervals. **(d)** Maps visualizing the average feature weights across participants, for each training day. On each map, channels are represented on the y-axis and individual frequency values on the x-axis. Scalp topographies below each map display the average weight for each channel, across all frequencies and participants, on each training day. **(e)** The bar plot represents statistical results (t values) when testing for changes in frequency contribution with training. Positive and negative value indicates respectively a decrease and increase in contribution throughout the training period. Yellow bars highlight the frequency(ies) for which the test was statistically significant (p-value < 0.05, permutation test). Scalp topographies show the results of the same statistical analysis performed on the scalp space, separately for two frequency intervals. These were defined based on the analysis in the frequency space, and include respectively the 2-10 Hz (low-frequency interval of interest) and 52-66 Hz (high-frequency interval of interest) range. A decrease in the contribution of a given electrode in the classification is indicated in blue, an increase in red (t values). Black dots highlight electrodes showing a statistically significant effect (p-value < 0.05, permutation test). **(f)** Schematics of the approach used to extract a global index to quantify the change across the 5 days of training. For each feature in the channel x frequency space, the euclidean distance between two consecutive training days is calculated, then the index is obtained by averaging the resulting 4 distances. **(g)** Euclidean distance index for each individual feature. **(h)** Correlation between the global index obtained with the euclidean distance approach and the average BCI performance across the 5 training days.

#### EEG data preprocessing and analysis

EEG data recorded during the *online* session were preprocessed using Fieldtrip^49^ and the Semi Automatic Selection of Independent Component Analysis (SASICA^50^) toolboxes within the MATLAB environment (version R2018b; The MathWorks, Natick, MA, USA). The data were first filtered using a zero phase butterworth bandpass filter with cutoff frequencies of 1 and 70 Hz. They were then divided into epochs of 12 seconds centered around the syllable imagery onset, including 4 seconds pre-stimulus (during the fixation cross and the cue presentation) and 8 seconds post-imagery onset (5 seconds of online BCI control and 3 seconds of rest). Noisy channels and epochs were removed via visual inspection, after which the data were re-referenced to the common average (which also served the purpose to retrieve data from the reference channel). Principal Component Analysis (PCA) was used to identify and remove ocular and muscular artifacts. The choice of the components to be removed was guided by different metrics such as autocorrelation, focal trial activity, and dipole fit residual, computed through the SASICA toolbox. Last, noisy channels that were initially removed were added back to the dataset by interpolation.

#### Power changes during BCI-control

To investigate oscillatory modulations associated with speech imagery and BCI control, we computed the power change for each individual frequency and each channel during the entire trial with respect to the baseline activity in the −3 to −2 seconds pre-imagery onset (i.e. during the fixation cross presentation). For this, we used Morlet wavelets to decompose the preprocessed EEG time series in the time-frequency domain. The power values were baseline-normalized at the single trial level and separately for each frequency band, expressing the change in percentage.

The statistical significance of the power modulation averaged across the 5 training days was assessed separately in several frequency band of interest (namely theta: 4-7 Hz, alpha: 8-13 Hz, beta: 14-26 Hz, low-gamma: 27-40 Hz and high-gamma 41-70 Hz) at the level of scalp topography by considering the average over the 5 seconds of real-time BCI-control. To do so, we used a Monte Carlo test with 1000 permutations, shuffling baseline and BCI-control labels, and a 2-tailed paired t test to compare the average power over the two intervals. We used a standard cluster-based correction to account for multiple comparisons over the scalp ^49^. To assess the statistical significance of the identified clusters, their size was compared to the distribution of cluster sizes expected under the null hypothesis. Clusters that had a p-value < 0.05 (two-tailed) were considered significant.

#### Changes in EEG power across training

To assess the evolution of the neural activity during BCI-control over the 5 training days, we considered the average power for each frequency band and tested for a linear increase/decrease using the same approach based on planned contrast as used for behavioral performance (*LMM with planned contrast*). For this, we first averaged the power over the 5 s of real-time BCI-control, separately for each channel, day, and participant. Next, a linear mixed model was fitted for each channel and frequency band separately. The statistical significance of the model was assessed using a permutation test at a standard alpha threshold of 0.05: for each participant, the data were randomly permuted across days and this procedure was repeated 1000 times. Multiple comparison correction was performed based on the same clustering method as mentioned in the previous section.

#### Evolution of Brain-Behaviour over 5 days of training

Next, we investigated how the relationship between the power modulation in each frequency band and BCI-control performance changes over the training period. To do so, we performed separately for each frequency band and channel, a linear mixed model with BCI performance as a dependent variable, the same planned contrast used previously to model a linear change across the 5 training days and EEG power as independent variables. Participants were considered as a random factor, leading to the following model: *BCI_performance* ∼*EEG_power*day + (1|Participant)*. Significance was calculated using permutation tests followed by cluster-based multiple comparison correction (see previous section). In particular, we considered the coefficients for the interaction term “*EEG_power*day*”: a positive coefficient would indicate that the effect of power on BCI-control performance becomes stronger with training, and vice versa.

#### Electromyography analysis

EMG was recorded throughout the entire experiment by means of two bipolar electrodes placed over the participant’s face to control for a potential contribution of muscular activity to the classification between the two syllables. We assessed this potential bias by computing the classifier’s features and CV accuracy considering exclusively data recorded from the two EMG electrodes, separately for the offline and online session. We then compared these CV accuracies, averaged across training days, with those obtained with the EEG data using a two-tailed t-test. Furthermore, we tested for a linear improvement in CV accuracy throughout training during the online session, using separately the EEG and EMG data (“*LMM with planned contrast*”).

Last, we evaluated the similarity between the feature’s weight frequency-wise extracted with the EMG-classifier and EEG-classifier: an overlap between the frequency profiles would likely indicate a strong muscular contamination in the discrimination between the two imagined syllables.

All statistical analyses were carried out using MATLAB (version R2018b, The MathWorks, Natick, MA, USA) and R version 4.1 (R Core Team, 2021).

## Results

### Training improves BCI-control abilities and decoding accuracy

First, we tested for a change in BCI-control performance throughout 5 training days. The average performance showed a significant linear increase from Day-1 to −5 (F_1,59_ = 5.92, p < 0.05, η^2^_p_ = 0.09, Fig.1c), indicating that imagined speech abilities can get better with training. The improvement was found in the majority of participants (11/15, Fig. 1b), with marked inter-individual differences. Interestingly, we found a positive correlation between the average BCI-control performance obtained considering the entire training period, and individual learning slopes (r = 0.55, p < 0.05, Fig. 1d), showing that the best performers were those who also benefited the most from training.

As expected, there was no difference in performance between the two syllables (T_14_ = 1.64, p > 0.05, d = 0.42). We found a trend towards significance when testing for differences in performance between the four blocks (F_3,42_ = 2.54, p = 0.07, η^2^_p_ = 0.15), due to lower performance during the second block as compared to the third (T_14_ = 2.2, p = 0.045, d = 0.57) and fourth (T_14_ = 2.2, p = 0.043, d = 0.57).

The linear trend for improved performance was confirmed by an increase in the cross-validation (CV) accuracy obtained when computing the classifier parameters on *offline* data (F_1,59_ = 9.35, p < 0.01, η^2^_p_ = 0.14, Fig. 1e-left, blue line). This measure considers exclusively neural data, as no feedback is provided during the first part of the experiment. Using the same post-processing approach, we computed the CV accuracy on the *online* data. We observed a significant improvement over the 5-days of training (F_1,59_ = 17.79, p < 0.001, η^2^_p_ = 0.23, Fig. 1e-left, red line). Across training days, the CV accuracy was significantly higher in the *online* than *offline session* (T_14_ = 8.3, p < 0.001, d = 2.14). This difference was not due to a difference in the learning dynamics, as we found no statistical difference between the learning slope in the offline and online sessions (T_14_ = 1.5, p > 0.05, d = 0.38, Fig. 1e-right). All participants except 2 received at least one monetary bonus, reflecting performance improvement from one day to the next.

### Analysis of the classifier’s features

In a second step we analysed jointly participants’ performance and the features used by the classifier to distinguish the imagery of the two syllables. We found a marked correlation between the individual BCI-control performance and the features weight (r = 0.83, p = 0.00013, Fig. 2a). This indicates, as expected, that participants performing better present more discriminant features.

We then investigated which frequency bands contribute most prominently to the syllable discrimination and found a peak in contribution straddling the alpha and the lowest end of the beta interval (8-16 Hz), as well as the gamma band (Fig. 2b). In particular, we observed a linear increase in the average feature weights throughout the entire gamma interval, with the highest values for the highest frequencies up to 70Hz. The topographical representation of the features according to their frequency shows that the first peak was associated to a cluster over the left central region (Fig. 2c-left), whereas the gamma band contribution originated from temporal regions, bilaterally, and posterior-occipital areas (Supplementary Figure 2, Fig. 2c-right). A third, distinctive spatial pattern was found in the theta band, and characterized by a strong contribution from frontal and temporal regions (Supplementary Figure 2).

#### Features evolution over training

First, we considered the global evolution of the features (sum of the first 200 features) and found a significant linear increase in the weights over the course of the training (F_1,59_ = 8.62, p < 0.01, η^2^_p_ = 0.13, Supplementary Fig. 1b).

Next, we qualitatively inspected the change in feature weights both in frequency and spatially across the scalp. We found that across participants the most discriminant features consistently localized over temporal regions and involved most prominently frequencies above 30 Hz on each training day (Fig. 2d). While the feature weight distribution on the first two training days was rather scattered over the whole frequency-channel space, with more training it narrowed down to the most discriminant feature clusters (Fig. 2d).

To quantify this change, we first investigated the weight evolution frequency-wise and found two frequency ranges that presented a linear change throughout the training period: while the contribution of the 2-10 Hz interval decreased, that of the 52-66 Hz interval increased with training (Fig. 2e).

To identify which regions underpinned this effect, we considered the average over each of these two intervals, and ran the same analysis at the individual electrode level. We found a decrease in low-frequency contribution over bilateral temporal and frontal regions, together with an increase in the high-gamma band over left fronto-temporal regions (Fig. 2e)

We then investigated the link between the change in BCI-control performance and the change in the feature space. We extracted, separately for each participant, a global index representing the amount of change both in the frequency and spatial domains, computed as the euclidean distance between the weight of two consecutive days (Fig. 2f). By visually inspecting the euclidean distance for each feature (averaged across participants), we observed a strong overlap of this index (Fig. 2g) and the frequency-channel feature maps (Fig. 2d). We found a strong correlation between the average BCI performance and euclidean distance (r = 0.85, p < 0.001, Fig. 2h), indicating that participants performing better during the real-time control of the visual feedback were also those whose features changed most during training.

### Power modulation during BCI-control and neural changes over training

We subsequently investigated neural changes occurring throughout the BCI-control training considering the power modulation in each frequency band during the 5-s of BCI-control (*online* session). We first inspected changes during real-time control with respect to the baseline activity (i.e. during the last second of the fixation cross presentation), and found a significant power decrease over frontal and left-central electrodes in the alpha band, and a similar but more widespread pattern in the beta range (Supplementary Fig. 3). We also found enhanced power over posterior-occipital electrodes in the high gamma band (Supplementary Fig. 3). Next, we investigated linear changes in power modulation throughout training. We found an overall power increase from day 1 to 5 on all frequency bands (Fig. 3a). In particular, theta and low-gamma bands showed the strongest and most widespread increase in power across training (Fig. 3a-b). Smaller clusters of power increase were also found in other frequency bands, namely alpha, beta and high-gamma bands.

**Figure 3:**
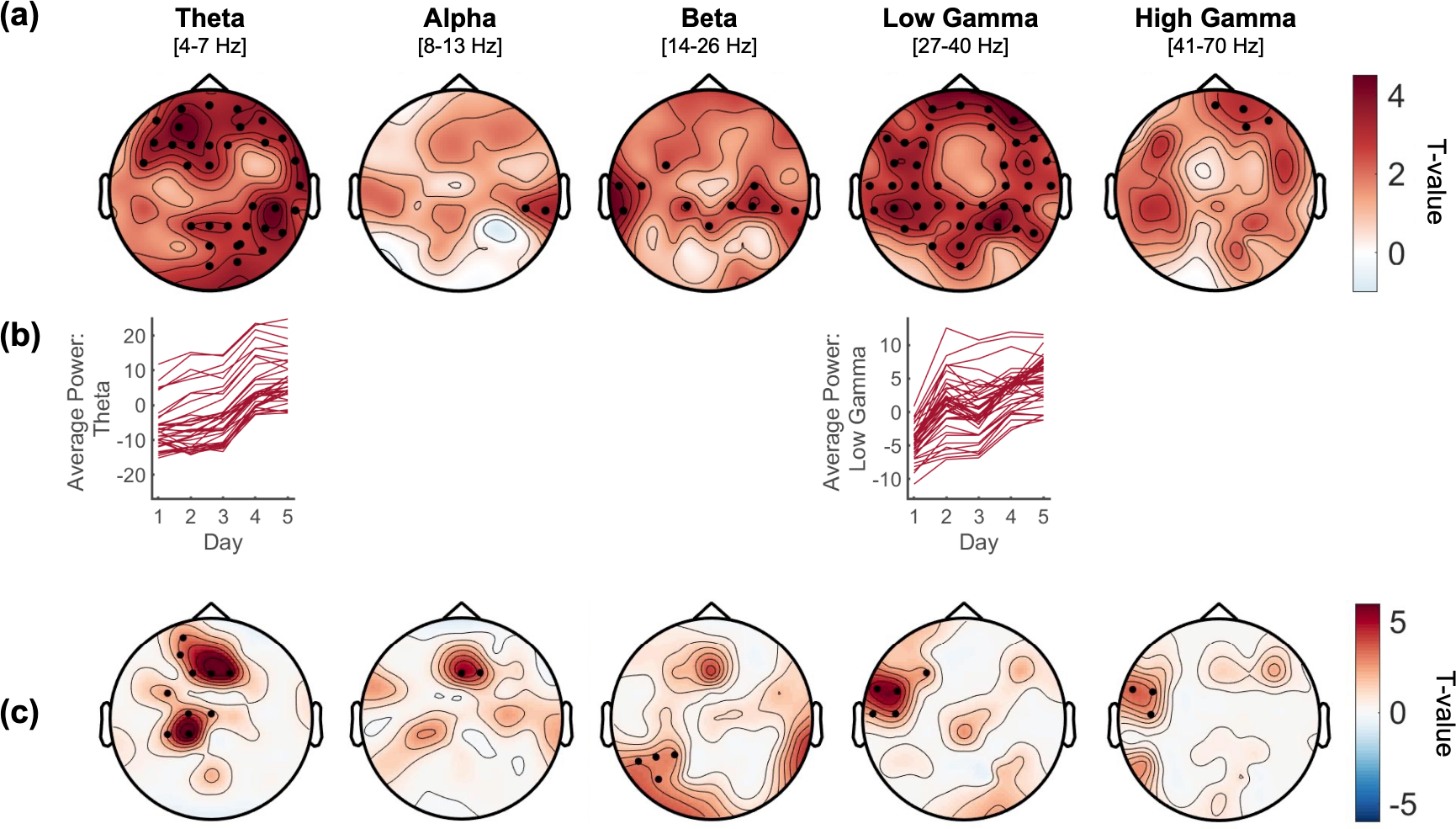
Evolution of power modulation over the 5 training days and relation with BCI-performance dynamics. **(a)** Topographies, one for each frequency band of interest, display the changes in EEG power across the 5 days of training. All frequency bands show a global linear power increase (positive t values). Electrodes showing significant linear trends are highlighted with black dots (permutation test, cluster corrected, p < 0.05). **(b)** Power (averaged over participants) for each day, plotted for each significant channel, shows a clear increase over the 5 training days. **(c)** Topographies, display the statistical result of the interaction term between EEG power and the effect of training (modelled with a planned contrast) on BCI-performance. Several statistically significant clusters display positive t-values, indicating that EEG power in specific regions and frequency bands became stronger with training. Among these, the most prominent were found over frontal and central regions in theta band and left temporal area in the low-gamma band. Significant electrodes are highlighted with black dots (permutation test, cluster corrected, p < 0.05).

### BCI-control performance and neural changes over training

Last, we investigated the link between BCI-control performance and power modulation over the 5 training days. We used a linear mixed model with BCI-performance as a dependent variable and, as predictors, the power in each frequency band and the planned contrast modelling a linear positive trend throughout training. The analysis revealed several clusters of electrodes showing a positive interaction between the two predictors indicating that power variations in these clusters increase their impact on BCI performance with training (Fig. 3c). In other words, specific regions and frequencies show dynamical changes in the direction of a stronger contribution in determining BCI-control. These included clusters located over frontal and central regions in the theta band, and over the left temporal region in the gamma band. We found additional smaller significant clusters over the central region in the alpha band, and the left posterior regions in the beta band.

### Comparison of syllable decoding from EEG and EMG

To ascertain the neural contribution to syllable decoding, we probed the potential contribution of EMG activity in discriminating between the two syllables. As we asked participants to imagine saying the syllables, a residual muscular activity cannot be excluded. We thus computed the CV accuracy of a model using exclusively EMG during the *offline* and *online* sessions. The average CV accuracy was above chance for both sessions. We then compared the EMG-based CV accuracy values to the EEG-based CV accuracy and found opposite effects for the two sessions. While CV accuracy for EMG data was higher than for EEG during the *offline* session (T_14_ = 2.2, p < 0.05, d = 0.57, Fig. 4a-left), the opposite was found for the *online* session with higher CV for EEG than EMG data (T_14_ = 2.77, p < 0.05, d = 0.71, Fig. 4a-right). Overall, CV accuracy based on online-EEG data was the highest (as compared to both EMG session and EEG-offline). Importantly, while EEG-online accuracy showed a strong linear increase throughout training (F_1,59_ = 17.79, p < 0.001, η^2^_p_ = 0.23), the same analysis performed with EMG data revealed no statistically significant change (F_1,59_ = 2.53, p > 0.05, η^2^_p_ = 0.04). Additionally, we found the distribution of the average weights obtained from the EMG classifier (Fig. 4b) to be markedly different from the histogram obtained considering the EEG features (Fig. 2b). Of note, the different magnitude in the range of the average weights between the EEG and EMG feature space in Fig. 2b and Fig. 4b is due to the lower number of EMG features (there were only two EMG channels versus 64 EEG channels). Although we cannot completely rule out the presence of subthreshold motor activity despite the explicit instruction not to perform articulatory movements, the EMG contribution remains marginal relative to the increased decoding accuracy over training.

**Figure 4:**
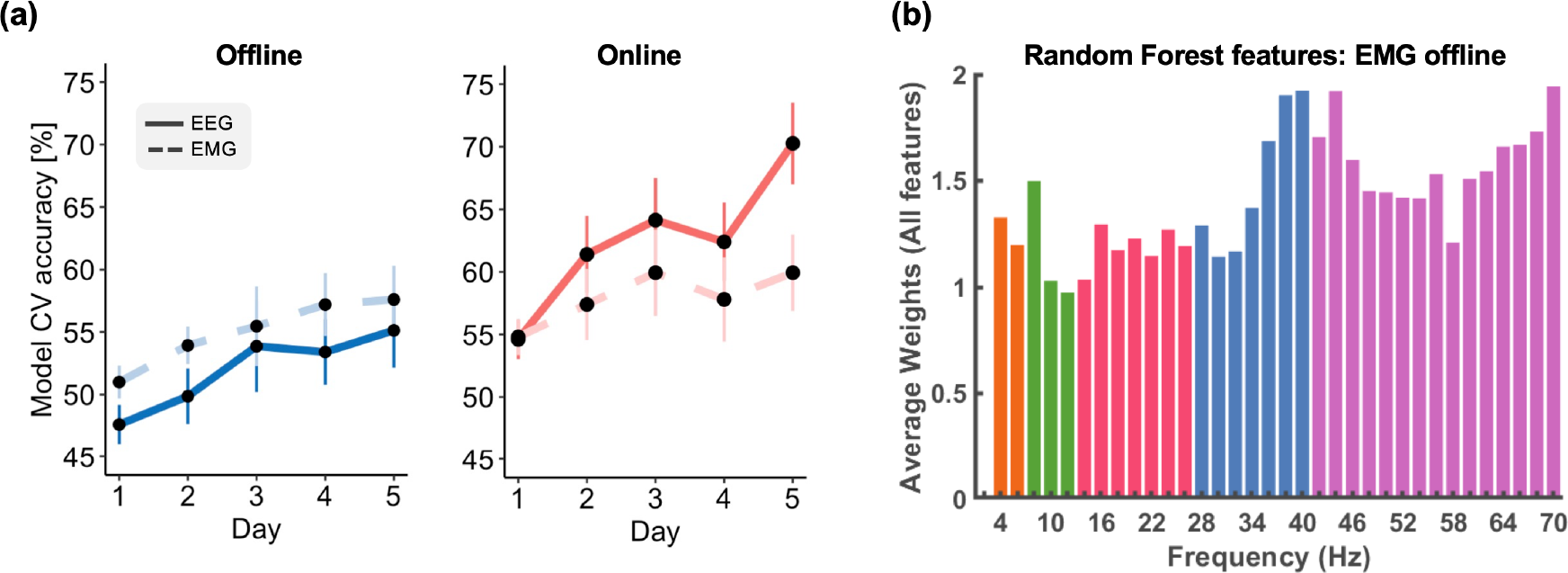
Decoding based on electromyographic (EMG) signals. To measure the contribution of potential muscular activity to syllable decoding, we computed a classifier based solely on EMGs from the right zygomaticus major and the orbicularis oris, separately for offline and online sessions. **(a)** CV accuracies based on EMG (dotted lines) and EEG (solid line) data acquired during the offline (blue) and online (red) sessions. **(b)** Histogram representing for each frequency, the average features’ weights obtained considering the EMG activity recorded during the offline session (the histogram is to be qualitatively compared with Fig. 2b).

## Discussion

We demonstrated that healthy individuals can learn to control an EEG speech-BCI by training over 5 consecutive days, and we uncover the neural mechanisms underpinning the acquisition of BCI-control skills. Learning to operate a BCI based on covert tasks has hitherto been investigated almost exclusively in the motor domain^51,52^. In the field of speech-BCIs, previous studies showed that it is possible to improve operating an intracranial BCI through attempted speech^3,4^. Yet, this question has not been explored for imagined speech BCI-control.

Behavioral performance in real-time BCI-control improved across training, concurrently with an increase from 55% to 70% in the CV accuracy obtained in post-processing on the same dataset (*online* session). This trend for a decoding improvement over the 5 training days was also observed on the neural activity recorded during syllable imagery without feedback (*offline* session), although on each day the CV accuracy was markedly lower than in the *online* session. This difference highlights the relevance of providing real-time feedback to the user to enhance the discriminability of neural patterns, and is in line with improved performance with a higher feedback rate as previously shown in non-human primates^53^. While this effect might reflect stronger arousal during real-time control, it more convincingly reflects that continuous visual feedback provides an error-driven strategy to improve BCI control.

While the majority of the participants (11 out of 15) improved performance, there were marked inter-individual differences both in control skills and in learning slope, extending to speech-based BCI-control the phenomenon of “BCI-illiteracy”, well-known in motor-imagery BCIs^37,38,54^. Poor BCI-operability seems to affect even more severely imagined speech, as on the first training day performance was below chance in most of our participants, with no outstanding performer. This is likely due to the relatively weak neural signals elicited by speech imagery^17,20^ and to the limited access to deeper speech brain regions with surface EEG, in sharp contrast with the lateralized, focal, and superficial pattern elicited by hand motor imagery, which make it relatively easy decodable with scalp recordings. Quite predictably, the decoding improvement was stronger in better performers. A dichotomy in learners versus non-learners over the volitional control of individual neurons in mnemonic structures has been previously reported for a single training session^55^. Here, we show that this difference is present also when training is carried over a longer period of time but it appears mitigated by repeating the task over multiple sessions. Individual factors likely play an important role in acquiring BCI skills, including cognitive, affective, and somatic aspects^28,56^, whose investigation extends beyond the present work.

Given that the task was a simple binary syllable classification, we controlled for purely motor strategies by also analyzing the EMG. Decoding improvement during real-time control was absent when CV accuracy was computed (in post-processing) from EMG signals alone, and unlike with EEG data no distinctive decoding features could be identified. Although we cannot rule out the presence of subthreshold motor activation given the above-chance decoding on some EMG datasets, the better EEG-based decoding accuracies and their improvement over days during the *online* session show that learning mechanisms rely upon changes at the central, rather than peripheral, nervous system level.

The behavioral improvement was accompanied by specific changes in the decoding features as well as on the EEG power modulation during the BCI-control. On average, the most discriminant features were located over the temporal regions bilaterally in the gamma band, and over the left sensorimotor cortex in the 8-16 Hz range, overlapping with key speech regions^39,57^ previously exploited as decoding sites^6,30^. However, over training we qualitatively observed a pruning effect, with the least discriminant features progressively decreasing their contribution in favor of more focal clusters around the features most contributing to the classification. The contribution of lower frequencies in frontal and temporal regions decreased in favor of a stronger involvement in high-gamma band in frontal and left centro-temporal regions. A similar parsimonious behavior is observed in fMRI-neurofeedback training, resulting in a reduction of redundant connections while strengthening the relevant ones in a restricted set of brain regions ^58^. Whether this effect is more pronounced when the action is performed in absence of any motor output, such as in an imagined speech BCI, has yet to be determined. Interestingly, we also found that a larger degree of change in the feature space was associated with higher BCI-control performance. This points to the importance of a dynamic calibration of the decoder parameters while the BCI is being operated, such as with adaptive classifiers that are able to account in real-time for learning-related changes^59^.

The power of the neural activity elicited by BCI control (irrespective to the imagined syllable) also substantially increased across training over the entire spectrum, most prominently in the theta and low-gamma band. Both frequency bands are highly relevant in speech perception and production, respectively underpinning syllabic and phonemic processing^60,61^. Interestingly, we found that the same two frequency bands showed more prominently clusters strengthening their influence of BCI performance as training progressed, specifically over fronto-central regions for theta power and over the left temporal area for the low-gamma band. The fact that the contribution of the theta band as a decoding feature decreased (Fig. 2e) while undergoing substantial power increase across training (Fig. 3a) in relation to performance improvement (Fig. 3c) might appear contradictory. In this context, however, the change in theta power might reflect mnemonic changes^62^ via changes in synaptic plasticity^63^, hence not directly contributing to syllable classification, even that occurring over the same time scale as the training duration in our experiment^64^. Further analyses are necessary to elucidate a potential top-down role of the theta band on higher frequency bands involved in discriminating between the two syllables.

## Conclusions

The present study fills an important gap in the field of imagined speech BCI suffering from low performance by showing that controllability can be improved with training, even when starting from chance-level performance. It provides solid neurophysiological grounds to improve current BCI systems based on speech-imagery, notably by enhancing decoding using a pre-defined distributed yet focal subset of brain regions over temporal and fronto-central areas, that we found to be implicated in both decoding and learning. Although surface EEG-based BCIs are very unlikely to become stand-alone communication devices, they will find valuable applications elsewhere in the field of neurotechnology for language disorders. First, providing ease of set-up and the improvement of current decoders, rehabilitative interventions based on closing the loop with a real-time feedback on imagery attempts could be used to boost the recruitment of residual neural patterns and promote neural plasticity. Second, training to perform real-time control over several sessions could be used as a benchmark to identify learners who would benefit most from intracortical BCIs. The future of speech-imagery BCIs holds promise for a variety of purposeful scenarios, particularly those that rely on human-machine co-adaptation.

## Data Availability

The data that support the findings of this study are available from the corresponding author upon reasonable request.

### Acknowledgement

We thank the Human Neuroscience Platform of the Fondation Campus Biotech and Shizhe Wu for technical advice. This study has been supported by the National Center of Competence in Research “Evolving Language”, Swiss National Science Foundation Agreement #51NF40_180888.

## Supplementary information

**Supplementary Fig. 1.**
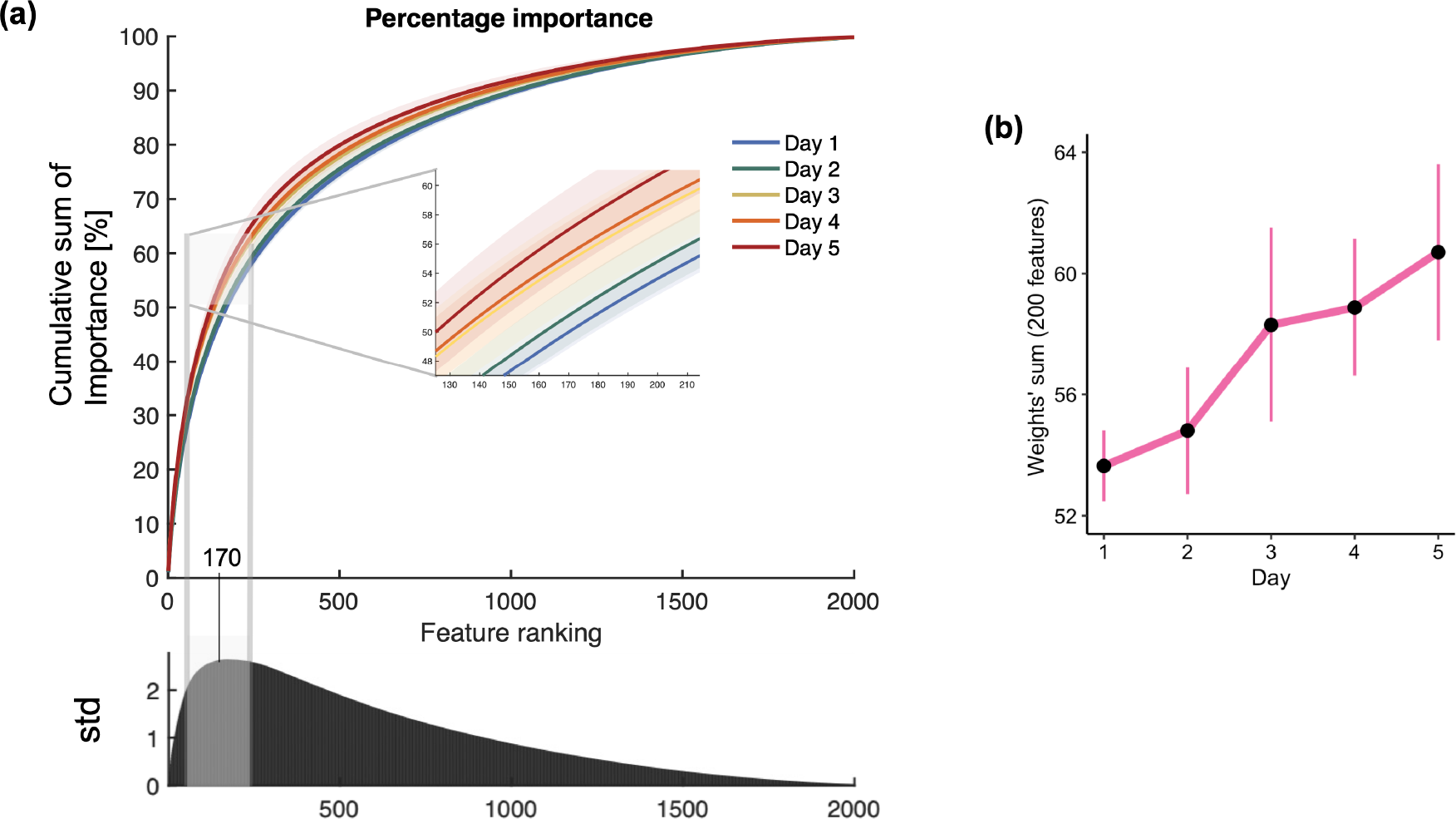
Analysis of the classifier’s features. The Random Forest classifier computes a weight for each feature (i.e. a channel-frequency pair), expressed as a percentage indicating how much the feature contributes to the accuracy of the model. The features are ranked according to their weight in descending order, from the highest to the lowest percentage. **(a)** Cumulative sum of the features weights along the ranking for each of the 5 days of training. The cumulative sum increases linearly from day 1 to day 5, so that as training progresses, a lower number of features is necessary to account for the same percentage. The value of 50% of cumulative importance is reached within the first 200 features across the 5 training days. The bottom plot illustrates, for each ranking position, the standard deviation calculated across the cumulative sums of the 5 training days. The variability increases up to the 170th ranking place, indicating that changes across training concentrate in a subset of most discriminant features and that features with lower ranking carry little information about changes over the 5 days. **(b)** Sum of the first 200 features’ weight across all participants across each training day shows a linear increase.

**Supplementary Fig. 2.**
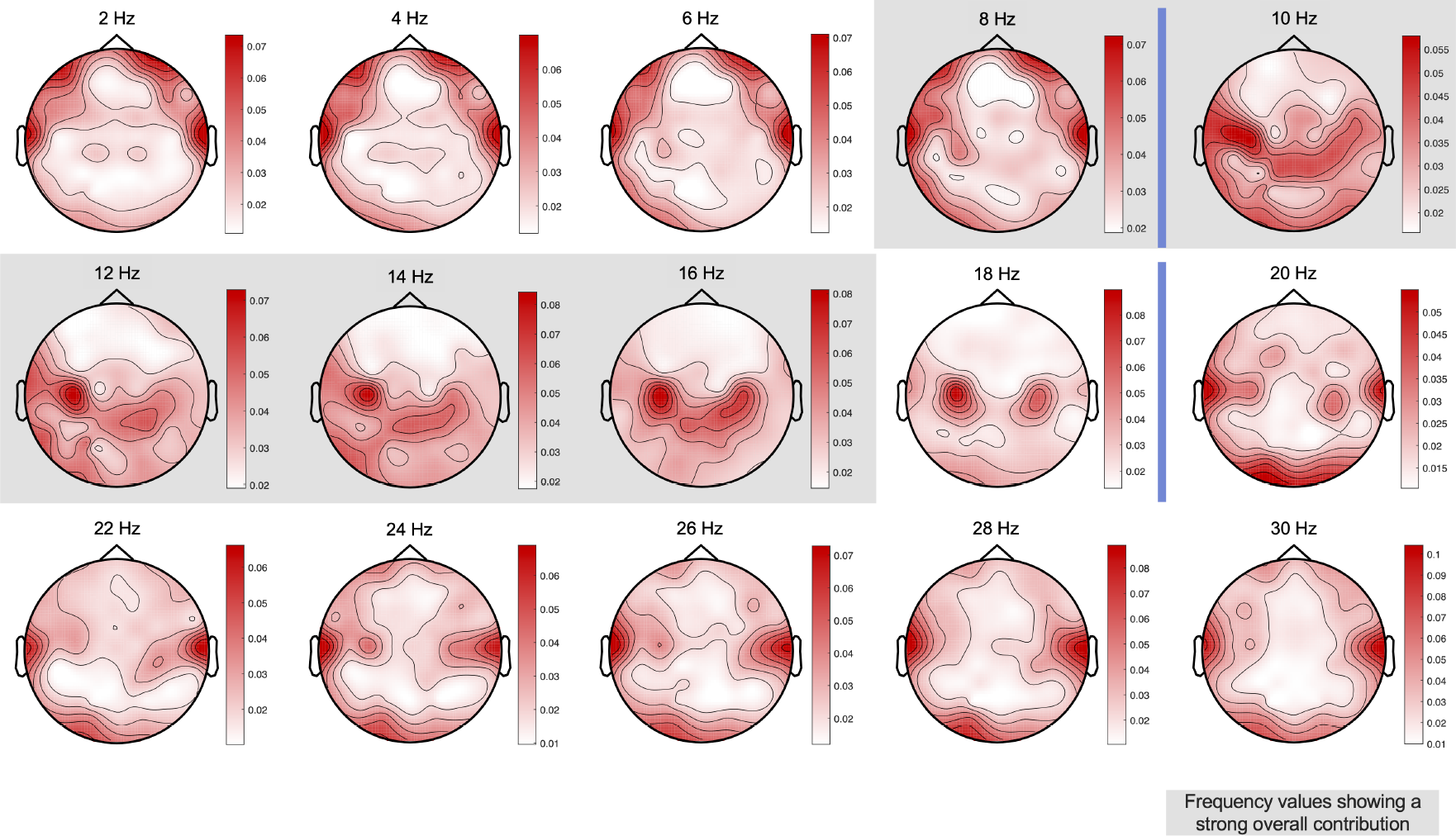
Topographies of features’ weight. Each topography represents the average features’ weight at a certain frequency, from 2 Hz to 30 Hz. Average values are obtained by considering data from all participants and the 5 days of training. Vertical blue bars delimit topographical transitions, whereas grey backgrounds indicate topographies within one frequency range contributing the most to the discrimination between the two syllables (8-16 Hz, see Fig. 2c). Topographies beyond 30 Hz are not shown as strongly overlapping with the last one, at 30 Hz.

**Supplementary Fig. 3:**
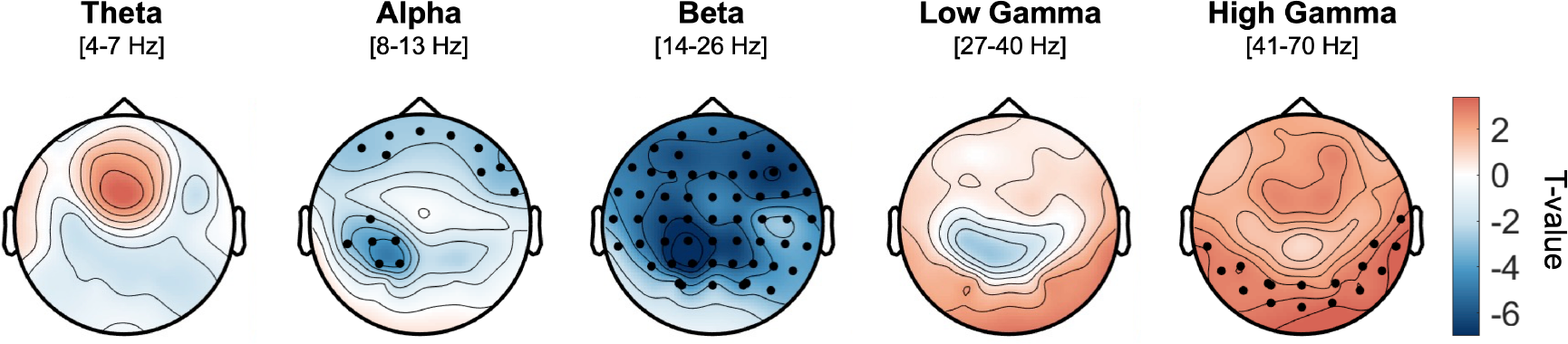
Power modulation during BCI-control. EEG power modulation in the overall dataset (all participants and days) relative to the baseline (last second of the fixation cross). Power data averaged between 0 and 5 s for each frequency band were compared against the baseline using a within participant paired t-test to find the channels showing significant power increase (Cluster-based correction, two-tailed, target threshold α= 0.05).

